# Signatures of sex ratio distortion in humans

**DOI:** 10.64898/2026.02.04.702084

**Authors:** James G. Baldwin-Brown, Sergiusz Wesolowski, Raquel Mae Zimmerman, Bennet Peterson, Martin Tristani-Firouzi, Edgar Havier Hernandez, Kenneth I. Aston, Mark Yandell, Nitin Phadnis

## Abstract

Segregation distortion, the disproportionate inheritance of selfish genetic elements, is an important evolutionary force. While many species carry distorters, it is not clear if humans do. Major limitations for detecting human distortion are the small size of human families and the lack of genetic markers in most subjects. Here, we present evidence of strong distortion in a large human pedigree. We analyzed pedigrees from the Utah Population Database and identified lineages with a high chance of carrying a distorter. In particular, we identified a family that preferentially produced male offspring at a 2:1 ratio. This pattern is consistent with a distorting *Y*-chromosome, a rarity in species with degenerate *Y*-chromosomes. The detection of such non-Mendelian inheritance patterns suggests that human genomes may harbor segregation distorters.

## Introduction

Segregation distorters are selfish genes that bias Mendelian segregation in their favor. They often do this by manipulating the production or function of gametes to outcompete those with the alternative homologous chromosomes (Sandler & Novitski 1957, Lyttle 1976, Lindholm et al. 2016). Across eukaryotes, signals of segregation distortion have been identified through skewed segregation ratios (Morgan et al. 1925, Sturtevant & Dobzhansky 1936, Hickey & Craig 1966, Turner & Perkins 1979, Jaenike 2001, Lyon 2003, Fishman et al. 2008, Matsuda et al. 2011). Sometimes, multiple distinct segregation distorters can exist even within single species (discussed in Kingan et al. 2010). In mice, the *t*-haplotype on chromosome 17 causes segregation distortion in males by sabotaging the swimming of sperm that carry the wild-type chromosome 17 (Winkler and Lindholm 2022, Swanepoel and Mueller 2024). A gamete-killing distorter is also known in mice: divergence in copy number of ampliconic genes (genes with multiple copies) *Slx* & *Sly* causes sex ratio distortion (Coquet et al. 2012, Baird et al. 2023, Arlt et al. 2025, Campbell and Heitzmann 2025). Similar ampliconic genes exist in humans as well (Kruger et al. 2019, Ye et al. 2018, Bhowmick & Takahata 2018, Lucotte et al. 2018, Vegesna et al. 2019). Humans present no obvious biological exception in terms of harboring distorters. Given the wide distribution of distorters across eukaryotes, including in mammalian systems, it is surprising that none have been definitively identified in humans.

Distorters in natural populations are usually found serendipitously when performing controlled crosses and measuring the transmission ratios of alleles in large numbers of offspring. Sex chromosome distorters are especially easy to detect because they produce progeny with skewed sex ratios. For example, *X*-linked distorters often destroy *Y*-bearing sperm, producing heavily female biased progeny (Jaenike 2001). Anecdotally, many people know a family where most children are the same sex. Unfortunately, even the largest human families are too small to reliably detect distorters with offspring counting in a single generation. Segregation distorters can also reside on autosomes but detection of these necessitates the use of autosomal genetic markers. Small family sizes, long lifespans, and ethical constraints make the usual detection methods impractical to apply at scale in humans.

Early studies of single loci report some evidence of distorted transmission ratios in humans (Evans et al. 1994, Chakraborty et al. 1996, Naumova et al. 1998, Eaves et al 1999, Girardet et al. 2000, Naumova et al. 2001). Later studies took a more agnostic approach to detecting transmission distortion in humans. These attempts generally used microarray genotyping data for one or two generations of transmission of alleles in families. The first large-scale attempt to specifically look for distortion in humans, based on the amount of shared genetic material among siblings, argued for large-scale, small-effect transmission distortion throughout the human population (Zöllner et al. 2004). This study spurred interest in the field and was followed by three landmark studies using either 60 individuals from HapMap or 5209 individuals from the Framingham heart panel (Frazer et al. 2007, Santos et al. 2009, Deng et al. 2009 Paterson et al. 2009). All three studies used the Transmission Disequilibrium Test (TDT) to identify loci with biased transmission ratios (Spielman et al. 1993). They each found some number of autosomal markers that show evidence of transmission distortion. Deng et al. found 1,205 outlier SNPs and 224 candidate genes, Paterson et al. found 8 outlier SNPs with their most conservative thresholding, and Santos et al. found one site that was convincingly distorting.

A comprehensive reanalysis of these studies, however, showed that most of these biased loci were either not significant at the genome-wide level or best explained by genotyping errors (Meyer et al. 2012). Meyer et al. found the eight conservative SNPs identified by Paterson et al. and the site identified by Santos et al. to be true outliers. Meyer et al. identified two key limitations in accurately detecting transmission distortion in pedigrees. First, even minor errors in genotyping calls produce pervasive false positive signals of transmission distortion. Second, the small sample sizes of most genotyped pedigrees limit the power to detect anything but the strongest signals of distortion. Thus, these approaches have low power to detect true positives and a high likelihood of generating false positive signals of transmission distortion. A large amount of genotyping data from a small number of individuals exacerbates these problems. Finally, studies of small numbers of subjects are unlikely to sample distorters that are uncommon in the population.

Here, inspired by Meyer et al.’s insights, we attempt to circumvent the two bottlenecks in detecting transmission distortion in human pedigrees: genotyping errors and small sample sizes. First, we focus only on sex chromosome transmission using the recorded sex of individuals in pedigrees. To the extent that the recorded sex of individuals accurately reflects their sex chromosome genotypes, our approach should side-step the pervasive false positives arising from genotyping errors. Second, we are able to increase the sample size in our study by an order of magnitude compared to previous studies. Because our approach only requires the recorded sex of an individual, we can now access and use information from individuals going back more than 10 generations where molecular genotype information is unavailable. To the degree that the pedigree data do not suffer systematic biases in recording males vs. females, the sexes of individuals should reflect the transmission of sex chromosomes.

Despite the *a priori* rarity of *Y*-chromosome distorters, we focus on understanding *Y*-biased inheritance patterns for several reasons. First, the patrilineal inheritance and expression of *Y*-chromosomes in every generation makes it possible to deterministically track the inheritance patterns of *Y*-chromosomes compared to *X*-chromosomes. Second, because human males are hemizygous, deleterious alleles on either the *Y*-chromosome or the *X*-chromosome will harm males and produce a bias toward female offspring, but not toward male offspring. This property of the sex chromosomes makes it hard to distinguish whether female bias is due to deleterious alleles or drivers, whereas male bias cannot be confused in this way. Third, because males determine the sex of offspring in humans, any observed anomalies are more likely to be the result of male gametogenesis. Because *Y*-chromosomes exclusively carry genes required for male fertility and sex determination, and other sources of bias such as viability differences are unlikely to explain the observed patterns. Fourth, our approach treats the SRY locus that determines sex as a dominant, visible marker that faithfully tracks the entire recombinationally inert *Y*-chromosome, with the exception of the pseudoautosomal region. Difficult to genotype regions that may nevertheless contribute to human segregation distortion, such as heterochromatin and satellite content, ampliconic genes, non-coding RNAs, etc can be tracked using our method. We report evidence for a consistently male-biased human family. This non-Mendelian inheritance suggests that human genomes may harbor segregation distorters.

## Results

To look for signatures of distortion in humans, we used a Bayesian algorithm to detect distorted families in pedigrees. We used only the recorded sexes of 76,445 individuals from the Utah Population Database (UPDB), which offers a generational depth dating back to the 1700s (Smith & Mineau 2021, Slattery & Kerber 1993) (Utah Population Database). This is an order of magnitude larger than the previous largest study (Paterson et al. 2009). The overall sex ratio of individuals across the dataset is nearly Mendelian, with 50.2% males. We used a probabilistic programming approach that uses Bayesian networks to detect systematic sex biases within families. Our program, Warp, lets us find families with patterns of larger-than-expected proportions of males or females (Sup. Fig. 1).

Warp assigns a likelihood of carrying a transmission distorter to each individual in the pedigree. In the first step of Warp, each individual is assigned a low initial likelihood of carrying a transmission distorter (1 in 100 here, but the process is robust to many parameters tested – see Methods). These initial likelihoods start out very low because they are based on only a single data point per individual. Starting from individuals at the bottom of the tree, Warp looks at the parents of each individual. Using the chain rule and Bayes’ theorem, it then updates the children’s likelihoods of carrying a transmission distorter based on their parent’s sexes and likelihoods (Fig. 1, Sup. Fig. 1, Appendix).

**Figure 1.**
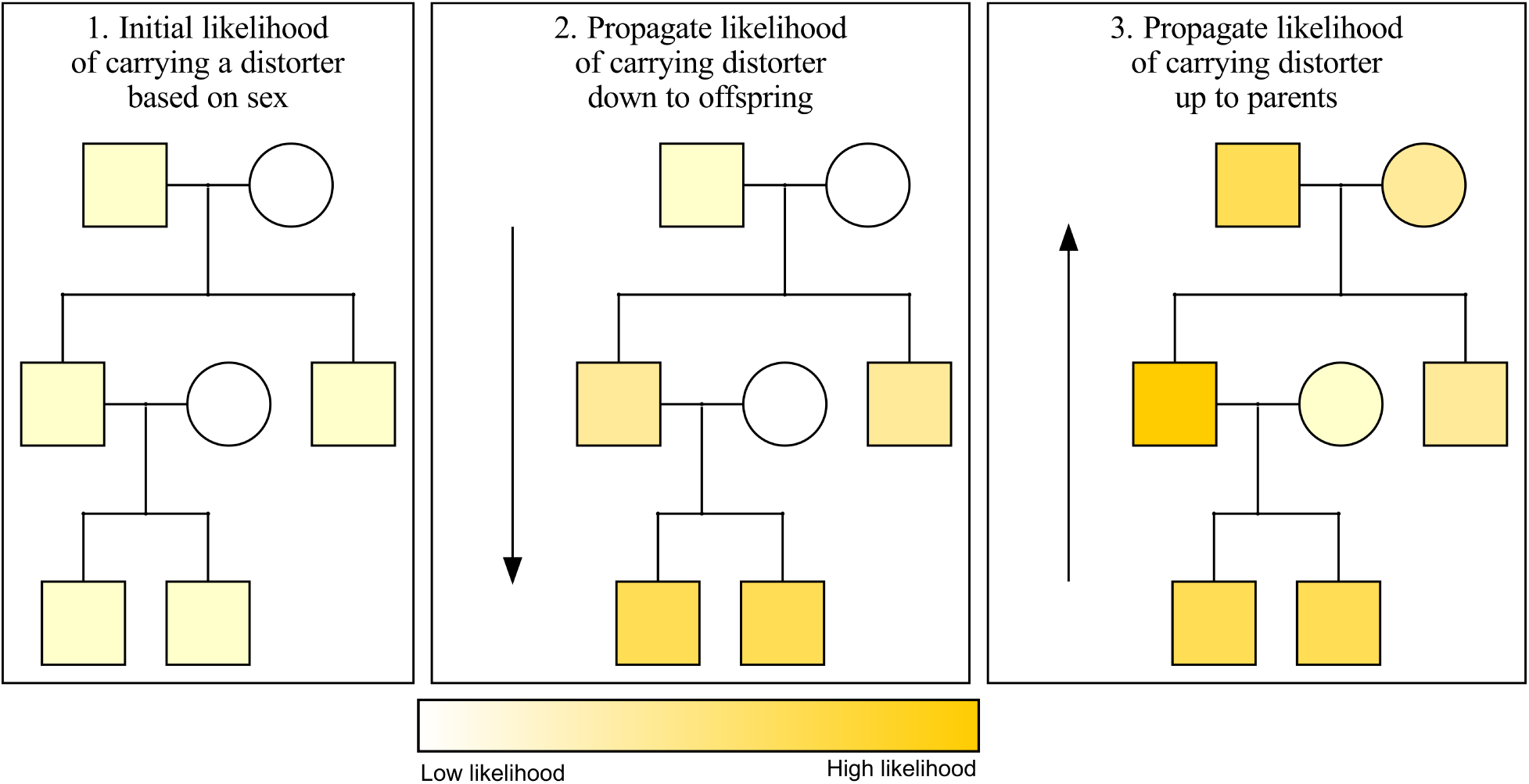
Warp has high power to infer the likelihood of carrying a distorter in pedigrees regardless of penetrance, inheritance pattern, and other challenges. Warp propagates the likelihood of carrying a distorter from parent to child, and from child to parent, using Bayesian network propagation (a combination of Bayes’ rule and the chain rule). Here, we show an abstract representation of the way Warp propagates probability information up and down a pedigree. Initial likelihoods for carrying the distorter are set using the phenotypes of each individual and the expected allele frequency of the distorter. In the second step, the likelihoods of all offspring of updated individuals are in turn updated based on the parent’s genotype onward down the pedigree. Finally, parents of updated offspring are also updated. This process is repeated until all likelihoods stop changing. In the case of non-cyclic pedigrees (those with no inbreeding), only one iteration is required.

Warp then iterates the same logic going down through the generations until it reaches the bottom of the pedigree. Once the bottom of the pedigree is reached, Warp iterates back up through the generations in the pedigree using the same logic. For example, a father of four sons has a higher likelihood of carrying a male-biased distorter compared to a father of two sons and two daughters. This process is iterated up and down through the pedigrees until the likelihoods stabilize.

Warp identified two putative male-biased families and six putative female-biased families in this dataset (Fig. 2a, Sup. Fig. 3, 4). Although we are motivated to identify signatures of sex chromosome distortion, viability differences can also bias sex ratios within families (Jaenike 2001). For example, the presence of an *X*-linked recessive lethal mutation in a family will produce fewer surviving males than females, generating a pattern indistinguishable from a female-biased distorter. In addition, tracking the transmission of each identical-by-descent *X*-chromosome is challenging. In contrast, viability differences are less likely to affect inferences about male-biased transmission. *Y*-chromosomes carry few if any viability-essential genes. Moreover, a *Y*-linked mutation that reduces viability would produce a female-biased family, and not male-biased. Male-biased families are therefore less likely to produce a spurious signal of transmission distortion. In addition, when searching for male biased distorters, we can easily track the transmission of each identical-by-descent *Y*-chromosome across many generations of patrilineal transmission in deep pedigrees. Together, because *Y*-chromosomes are hemizygous in males, largely non-recombining, not essential for viability (c.f. Turner’s syndrome, Ranke and Saenger 2001), and tracked without skipping generations, we focus our further analyses on male-biased families (Fig. 3).

**Figure 2.**
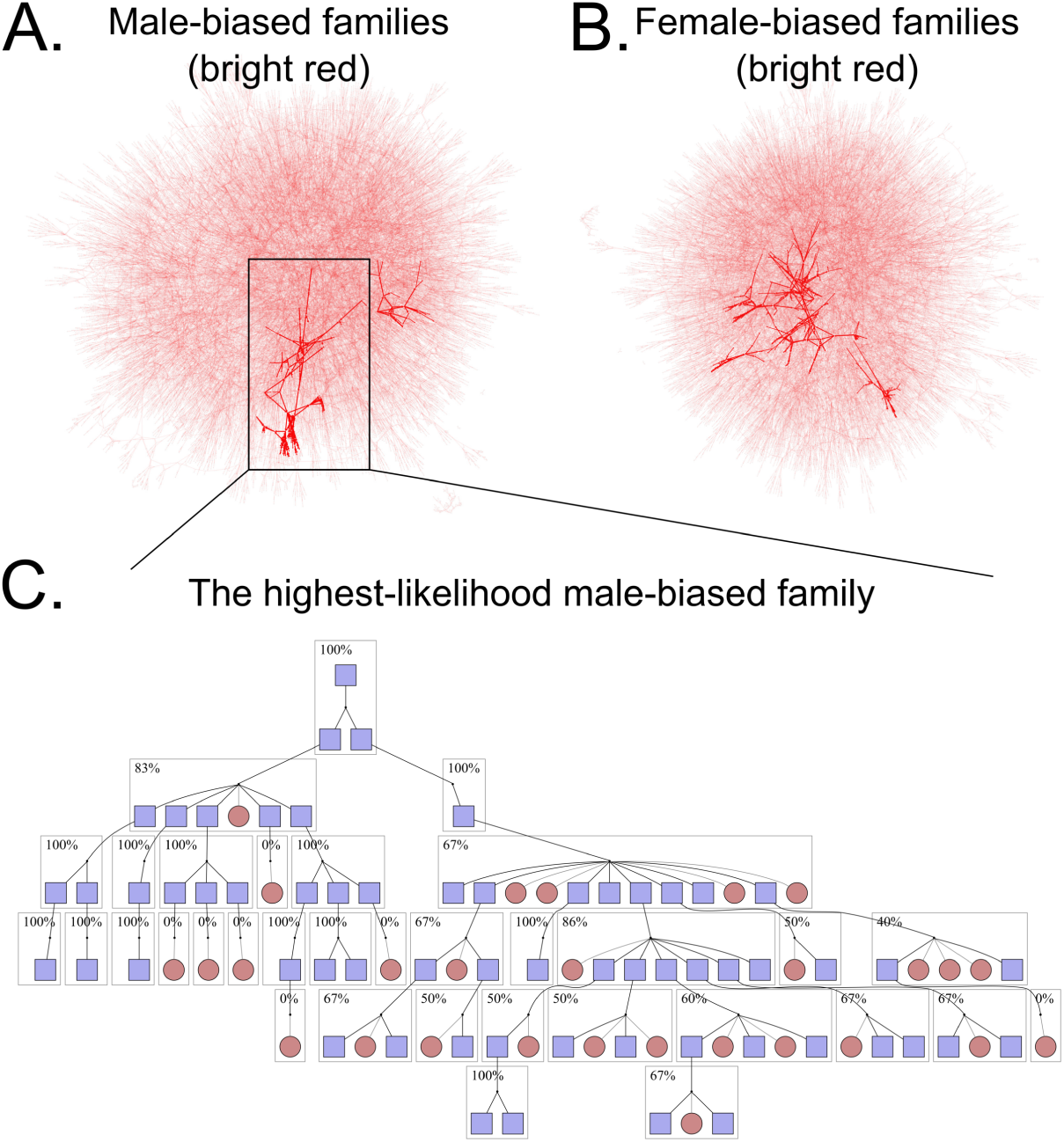
One human family shows a strong bias toward male offspring. A. Results from Warp testing for male bias plotted as a network of parent-child relationships (analogous to a pedigree). Points represent individuals and arrows represent parent-child connections. Bright red connections represent families that contain high-likelihood individuals. Two clusters are highlighted in red, indicating two potential distorters detected here. B. The same, but here testing for X-biased distorters. Six families are highlighted. C. A close look at the most significant family. A single male, top, carries a putatively distorting Y-chromosome such that about ⅔ of offspring are male. This partial pedigree shows the offspring of the focal male. Dark lines represent lines of inheritance of the distorting Y-chromosome, while light lines represent other lines of inheritance. Blue boxes represent males, while red circles are females. Dotted borders indicate individuals with non-informative offspring that are not shown. Boxed individuals are offspring of a distorter-carrying male and are included in the transmission disequilibrium test.

**Figure 3.**
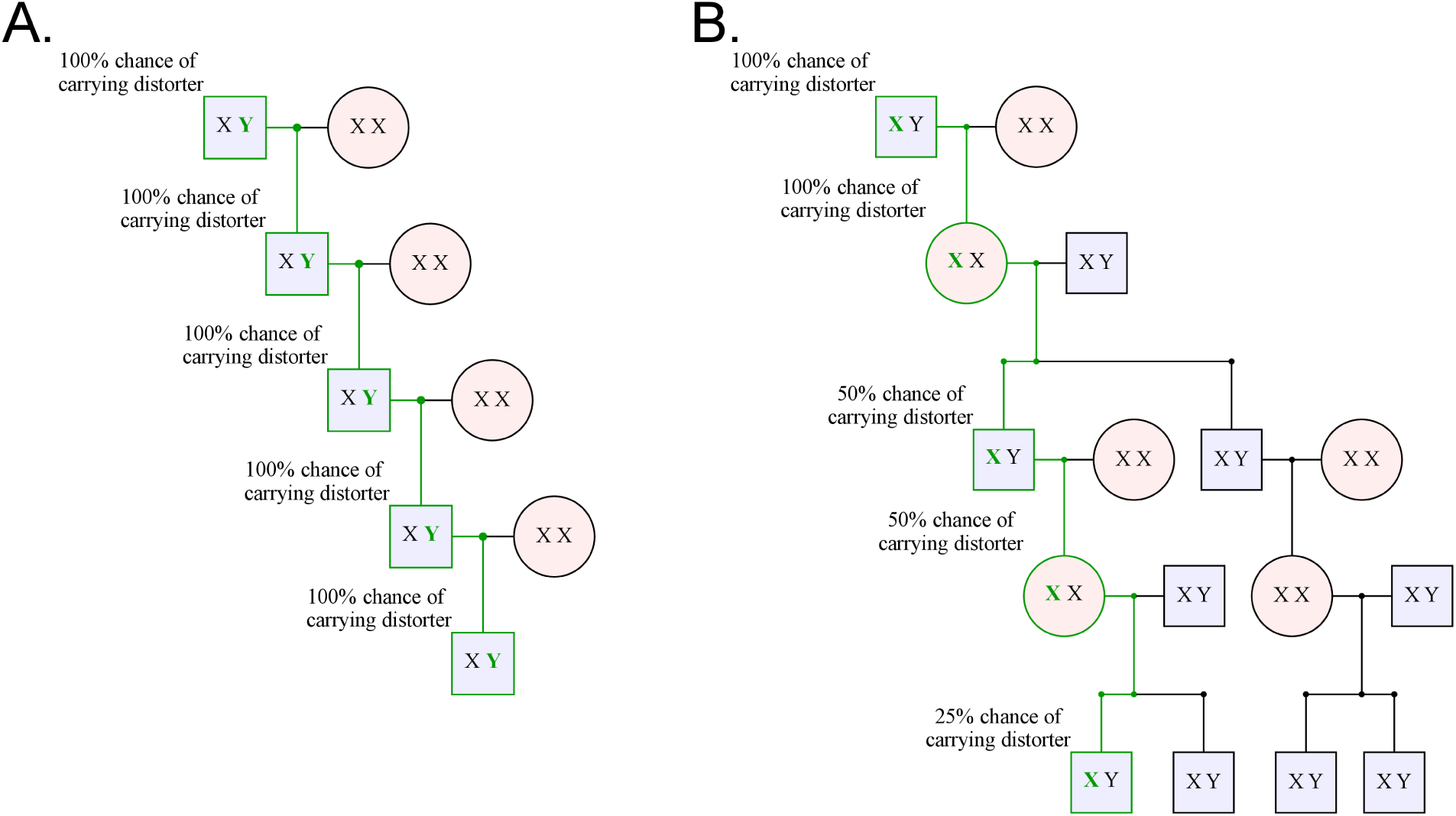
Y-linked distorters are more easily detected than X-linked distorters because of the simpler Y-chromosome inheritance pattern. A. A selfish Y-chromosome will always be inherited through the patrilineal line, allowing it to distort every generation. This makes finding all carriers in a patrilineal line easy: select for males. Here the bold green Y represents a Y-chromosome carrying a distorter, and the green line represents the line of descent of the distorter. B. On the other hand, a sex distorting X-chromosome only distorts in males, producing mostly females. It can, thus, only distort every other generation. To follow the distorter through the pedigree, we must also follow offspring that are not distorter carriers, “diluting” the family tree with non-distorting X-chromosomes that reduce the power of statistics like the TDT to detect distortion.

Although we detected some male- or female-biased families, any distribution, if sampled enough times, will produce some extreme outliers by chance. To test whether the dense clustering of males in outlier families occurred by chance, we compared the likelihood of observing this bias in our real pedigree to the likelihood of observing a similar bias in permuted pedigrees. To produce a conservative control dataset that most closely modeled the true data, we performed 1000 permutations of the whole pedigree dataset in which we randomly assigned sexes to individuals but kept the tree topology the same. To make the test as conservative as possible, our permuted pedigrees include extreme outcomes such as two individuals of the same sex having children. Comparing the likelihood values of individuals in the true pedigree to those of the permuted pedigrees showed that the high likelihood values of the most male-biased family in the true pedigree could not be explained by chance alone (*p* = 0.05 by highest-likelihood comparison, *p* = 0.017 by *z*-value comparison, and *p* = 0.001 by rank comparison, see methods). As an aside, we performed the same test on female-biased families and found a similar result (*p* = 0.004 by highest likelihood comparison, *p* = 0.17 by *z*-value comparison, and *p* = 0.017 by rank comparison). Together, these results provide the first indication that the sex biased families in our dataset would not occur by chance alone.

To complement our Bayesian approach, we next used the transmission distortion test (TDT), which is a workhorse test for detecting transmission biases in pedigrees (Spielman et al. 1993). The TDT is a chi-squared test that examines whether allelic transmission deviates from a 50:50 expectation. We identified all unique *Y*-chromosome lineages in the dataset and applied the TDT to each one. It is important to note that some “unique” *Y*-chromosomes may be identical by descent, but this shared ancestry may not be captured in the existing pedigree data. We then used false discovery rate (FDR)-corrected *p*-values from these results to see how many families were significantly male-biased. Because we do not trust outcomes from small families, we only include families with more than 75 assayable offspring in this analysis (an arbitrary cutoff). After FDR correction, only one patrilineal lineage out of 26,865 tested was significantly male biased (*p* = 0.0249, non-FDR-corrected *p* = 0.00102, Figures 2b, 4,). This patrilineal lineage identified by TDT is the same family that Warp indicated as most likely to carry a male-biased distorter. Because both Warp and TDT independently identify the same male-biased family, this family is likely to be a true outlier.

**Figure 4.**
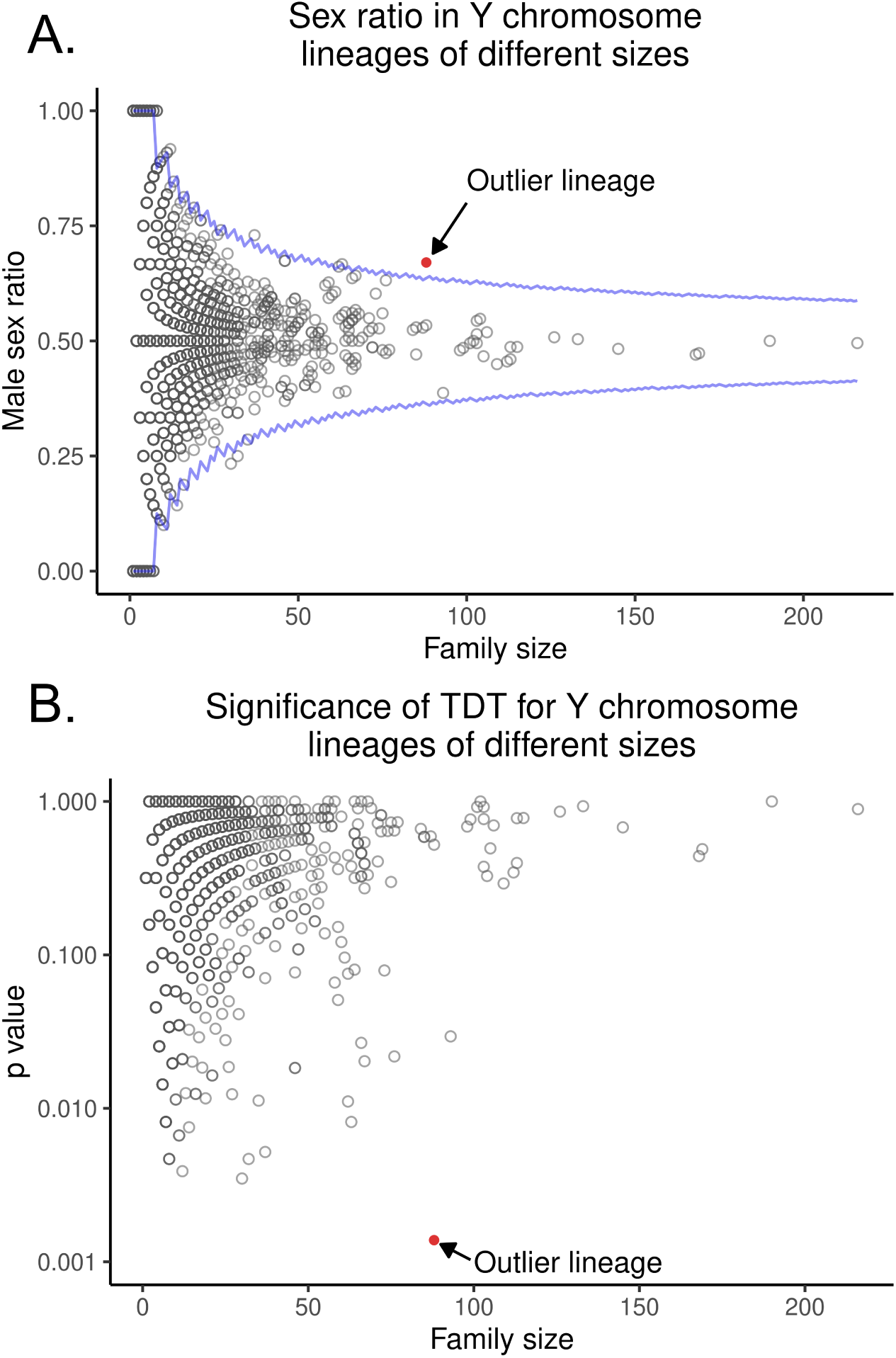
The TDT re-identifies the outlier male-biased family found be Warp. A. This figure shows the distribution in the UPDB of male sex proportion (y axis) in each Y-chromosome lineage compared to lineage size (x axis). Lineages with few individuals can have very distorted male proportions, but no lineage with a large number of individuals is as distorted as the focal family. The blue line is the theoretical 99% confidence interval. The red dot is the focal family discussed in the text. B. **The likely-distorting family is large and highly significantly distorted.** This figure shows the distribution in the UPDB of p-values derived from the transmission disequilibrium test. Each dot represents a unique patrilineal line of descent (and thus a unique Y-chromosome). The focal lineage discussed in the paper is red. The X axis is the number of children in each patrilineal line that contributed to the TDT test, and the Y axis is the p-value indicating the chance that the patrilineal line had such a high degree of sex distortion by chance.

The progenitor male of this family had two sons, both of whom produced highly male-skewed descendants. The first son had five male offspring out of six. While the other son produced only one son, this grandson produced eight male offspring out of 11. The patrilineal descendants of these sons also produced highly distorted sex ratios among their offspring. Together, this patrilineal line spanning seven generations had 33 males that carried the same *Y-*chromosome and produced at least one child, giving us high statistical power to detect systematic male bias. This family contained 89 informative transmissions with a total of 60 male offspring and only 29 female offspring. Put another way, 89 children contributed to the sex ratio of offspring; of those, 60 were male. This gives a final proportion of 67.4% male offspring in this family (Figure 4).

We next used Monte Carlo simulations that compare the observed sex ratio in this outlier family to 10,000 simulated families of equal size where the sexes of individuals were randomly assigned. This approach should be a numerical approximation of running the TDT on the family of interest. Here again, we allowed two individuals of the same sex to have children to include extreme outcomes. We compared the sex ratios of these simulated families to the real outlier family and found that the male bias in this outlier family was unlikely to occur by chance alone (We compared the sex ratios of these simulated families to the real outlier family and found that the male bias in this outlier family was unlikely to occur by chance alone (*p*=0.00138), just as the TDT indicated. Taken together, our results are consistent with the presence of a male-biased distorter in humans.

In summary, we used a combination of statistical techniques to first identify distorters, then confirm that they could not have arisen by chance. We used Warp to find high-likelihood distorter carriers, then used a permutation test in which we permuted the entire pedigree dataset to determine the chance of finding such a high-likelihood distorter. This produced several likely distorters and showed that they are true outliers from expectations. We then took the most likely *Y*-chromosome distorter and tested its entire lineage for distortion using the TDT. We then performed a follow-up Monte Carlo test within this lineage that agreed with the TDT. All of these tests together confirmed that this *Y*-chromosome is a likely sex distorter.

## Discussion

Sex ratio in humans has fascinated people for centuries (Bernoulli 1713, Fisher 1958, Graffelman and Hoekstra 2000, Sen 2003). Many researchers have shown that the human sex ratio is very close to 50:50, except in extreme cases such as war or other external pressures on sex-specific birth rates (Fisher 1958, Graffelman and Hoekstra 2000, Sen 2003). Based on our new results, we argue that comprehensive studies are likely to reveal the presence of selfish genes in human populations, including those that produce distorted sex ratios in some families. Identifying such selfish chromosomes in humans has many important implications. For example, given the dramatic effect on fertility observed in many known distorters in other species (Wu 1983, Jaenike 2001), distorters may explain the surprisingly high levels of male infertility in humans (Sharlip et al. 2002, Agarwal et al. 2015). Distorters may also help to explain long-standing questions related to human natural history. For example, certain regions of the Neanderthal genome have introgressed into the human genome, while other introgression ‘deserts’ are free of Neanderthal ancestry (Sankararaman et al. 2014). Strong selection preferring the Homo sapiens genome at these loci, such as that produced by distorters, could explain why these deserts remain Neanderthal-free.

Empirical studies of sex chromosome drivers in *Drosophila*, mosquitoes, and rodents show that *X*-chromosomes are more likely to harbor distorters than *Y*-chromosomes (Burt and Trivers 2009, Helleu et al. 2015). There are few known cases of *Y*-chromosome distortion, and they mostly occur in species with non-degenerate *Y*-chromosomes such as mosquitoes (Bachtrog 2013, Sweeny & Barr 1978). The human *Y*-chromosome is highly degenerate, with approximately 70 functioning genes (Jobling & Tyler-Smith 2017). None of these genes are known to be essential for viability, and typical XX females lacking *Y*-chromosomes are viable. A signal of *Y*-biased transmission distortion in humans is thus a surprise. Such *Y*-distortion, however, is known to occur in mammals. In mice, both experimental manipulations and studies in natural populations have shown *X*- and *Y*-chromosome distortion mediated by copy number variation in ampliconic genes, such as those in the *Slx*/*Sly* system (Coquet et al. 2012, Baird et al 2023, Arlt et al. 2025, Campbell and Heitzmann 2025). While humans lack *Slx*/*Sly* genes, they have other ampliconic genes such as the *RBMY* and *PRY* gene clusters that are candidates for playing a role in sex ratio distortion (Kruger et al. 2019, Ye et al. 2018, Bhowmick & Takahata 2018, Lucotte et al. 2018, Vegesna et al. 2019). The *PRY* gene cluster appears promising because it is involved in sperm elimination, a process recently shown to be a consequence of selfish chromosomes in the male germline (Ridges et al. 2026).

Although *Y*-chromosome distortion is a parsimonious explanation for the observed male-biased family, the exact molecular stage where biases such as *Y*-distortion occur in humans remains unknown. There are at least two known mechanisms underlying male gametic distortion. First, the most frequently observed mechanism for segregation distortion involves killing gametes bearing their competing homologous chromosomes (Jaenike 2001). Second, some selfish chromosomes, such as the *t*-haplotype on chromosome 17 in mouse, can distort transmission ratios by reducing the swimming ability of sperm that carry competing homologous chromosomes (Winkler and Lindholm 2022). Understanding the molecular and cytological basis of *Y*-chromosome distortion in our focal family would necessitate ethically deanonymizing affected individuals and following up with detailed cytological and genomic studies on sperm samples from individuals identified from the pedigree.

Although we focus on male biased families to detect signatures of *Y*-chromosome segregation distortion, we also found six putative female-biased families in our permutation tests. However, as discussed in the results, it is difficult to track *X*-chromosome inheritance patterns in these pedigrees without genotype information. Further, *X*-linked deleterious mutations can lead to a depletion of males in families, generating the same pattern as expected from *X*-chromosome segregation distortion. Without further information we are therefore cautious in interpreting the observed female biased families as due to the presence of an *X*-chromosome segregation distorter. Even more glaring, our genotype-free analysis does not extend to autosomal distorters where most human genomic material resides. This leaves open the possibility that human genomes harbor segregation distorters whose effects may depend on whether the genetic background is permissive for distortion or if it carries suppressor alleles.

Our work adds to several lines of evidence that point toward the presence of distorters in humans. First, previous attempts at large-scale searches for distortion have produced at least a few high-confidence outliers (Meyer et al. 2012). Second, the presence of ancient haplotypes on the human *X*-chromosome and the rapid expansion of the FT haplotype of the human *Y*-chromosome are suggestive of selfish transmission dynamics in human populations (Skov, et al, 2023; Hallast et al, 2020). Third, in an indirect method of detecting skews in sex chromosome transmission, a recent study argued for strong clustering of sexes within families, indicating the presence of heritable variation in sex ratios in humans (Wang et al. 2025; though see Zietsch et al. 2020). Together with our observation of apparent *Y*-biased distortion and the possibility of unexplored population-specific or suppressed distorters, segregation distorters appear likely to exist in human populations.

## Acknowledgements

Thanks to Michael Shapiro for his help revising the manuscript. We acknowledge NIH grants R01GM141422 and R35GM156267 to NP for funding this research. Thanks also to Molly Przeworski for her advice on the manuscript.

## Author contributions

Conceptualization, J.G.B., N.P., K.I., M.Y.; Methodology, J.G.B., N.P., K.I., M.Y., and RZ, SW, BP; Software, J.G.B. and RZ, SW, BP; Validation, J.G.B.; Formal analysis, J.G.B. and RZ, SW, BP; Investigation, J.G.B., K.I., and RZ, SW, BP; Resources, N.P., K.I., and M.Y.; Data curation, J.G.B.; Writing – original draft, J.G.B.; Writing – review & editing, J.G.B., N.P., K.I., M.Y., and RZ, SW, BP; Visualization, J.G.B.; Supervision, N.P. and M.Y.; Project administration, J.G.B., N.P., and M.Y.; Funding acquisition N.P. and M.Y

## STAR methods section

## Resource Availability

## Lead contact

Further information and requests for resources and reagents should be directed to and will be fulfilled by the lead contacts, James Baldwin-Brown and Nitin Phadnis.

## Materials availability

This study did not generate new unique reagents.

## Data and code availability

All data generated from this experiment will be made available to the scientific community and the public where ethically possible. All analysis scripts are available through GitHub at https://github.com/jgbaldwinbrown/jgbutils.

## Experimental model and study participant details

### Human participants

All analysis of pedigrees was based on fully anonymized pedigree data made available by the Utah Population Database.

## Method details

### Data generation

All data used here was derived from the UPDB and was collected previously; we have no new data to report (Smith & Mineau 2021).

## Quantification and statistical Analysis

### Bayesian detection of distorting pedigrees

The power to detect distortion is determined by pedigree size, distortion depth (as in how many generations the distortion is observed), breadth (how many distorted individuals per generation are observed), and the strength of distortion. In addition to those criteria, the observed distortion must adhere to Mendelian inheritance rules rather than being abundantly, but still randomly scattered throughout the family. To maximize the detection power and plausibility, we built a machine learning model that incorporates these three factors and contrasts the observed distribution of the distortion across family branches against a null model of the possibility of non-heritable causes.

Our model is based on the Probabilistic Programming paradigm and uses Bayesian networks (Stephenson 2000). In this model, the family pedigree structure is used as the backbone of the belief propagation algorithm (Kim et al. 1983). The belief propagation algorithm takes as input the observed phenotype (distorted or not) and updates the probability distribution for an individual, where the probability distribution says how likely the individual is to carry an allele that causes the observed phenotype. This is a simple application of Bayes’ Theorem, where P(Distorter) is updated to P(Distorter | Sex bias). That said, a single update is rarely enough to draw conclusions about the genetics of distortion, especially in the case of distorters with low penetrance or high prevalence. Thus, our algorithm extends the inference context to the whole pedigree. In the next step in the algorithm, each individual propagates the updated belief to its parents and children. This Bayesian Update (message passing) is conducted throughout the entire pedigree and propagated across generations, meaning that information from family founders can travel to the last offspring and vice versa. Each individual propagates and updates their beliefs until the algorithm converges and propagated beliefs no longer change the individual’s distribution – in other words, all individuals in the family agree as to whether their observed phenotype is due to heritable distortion or otherwise. In the process of reaching a consensus, the two models, coded in the conditional probability tables of the network, are competing to be the most plausible explanation. The null model assumes that the observed phenotypes are not due to heritability. When sex is the phenotype of interest, the model parameters are 50% sex penetrance, no heritability. The genetic distortion model assumes 90% penetrance and incorporates the probability of introducing or losing the mutation responsible for the distortion (de novo mutation rate 0.000001) and the a priori probability of a mutation occurring in the population (allele frequency 0.01). We chose this parameter combination by running Warp on all permutations of the following parameters, then selecting the parameter combination that produced the individual with the highest probability of carrying a distorter. Although this could cause cherry picking in another context, here we use Warp only to identify the most-likely-distorting families; the permutation tests of Warp, explained below, are the actual test of presence of distortion in the dataset. The permutations below are done with the same parameters for the true dataset and the shuffled dataset, so the parameters do not affect the significance of the permutation test. The tested penetrances were 50%, 60%, 62%, 64%, 66%, 68%, 70%, 75%, 80%, 85%, 90%, 95%, 97%, 99%. The tested allele frequencies were 0.001, 0.0001, 0.00001, 0.000001. The tested de novo mutation rates were 0.01, 0.001, 0.0001, 0.00001, and 0.000001.

We ran Warp on a pedigree reconstructed from 76,445 individuals for the UPDB. In such a large pedigree, by sheer chance, there are branches with sex bias present, but rarely can such biases be explained with heritable patterns.

Our statistical approach was designed to identify individuals that might carry a distorter, and was not designed with hypothesis testing in mind; therefore, we cannot assign a p-value to the hypothesis that a pedigree carries a distorter based on the core algorithm. To properly assign a p-value to this probability, we used a permutation test to calculate the FDR-corrected probability (p) that a pedigree could, by chance, contain the observed degree of distortion. This code is available at github.com/jgbaldwinbrown/tdt. We permuted the sex of all individuals in the pedigree 1,000 times and then re-ran Warp on each of these permuted samples. Each run of Warp on permuted data represents the outcome of a new pedigree of the same size, topology, and overall sex ratio, but no true evidence of distortion. We then compared the likelihood of being a distorter carrier in the most-affected individual in each dataset. The reported p-value is the percentage of permuted pedigrees that contain an individual with a higher likelihood of carrying a distorter than the individual in the true dataset with the highest likelihood. We also calculated the p-value in two other ways:

1. Within each run of Warp, we normalized all individuals’ likelihoods of carrying a distorter into Z values by subtracting the mean and dividing by the standard deviation. Then, we compared the Z value of the highest-likelihood individual in the true pedigree to the Z value of the highest-likelihood individual in each permuted pedigree. The percentage of permuted pedigree Z values higher than the true pedigree Z value is the p-value. This test resembles the above, but takes into account the fact that, for reasons we may not have considered, the mean and standard deviation of likelihoods in permuted pedigrees may differ from the true pedigree. A low p-value by this method indicates that the highest-likelihood individual is a larger outlier from all likelihoods in the true population than in the permuted populations.
2. We took the putative originator of the distorter in the true dataset and calculated its rank order position in the true population. We then compared this rank order position to the rank order position of the exact same individual in the permuted populations. The p-value here is the percentage of permuted populations in which this individual’s likelihood is higher-ranked in the permuted population compared to the true population. This comparison tests if the true likelihood of the most likely distorter is higher than expected in a randomized family, but does not take multiple testing into account the way that the above tests do. A low p-value here indicates that the putative originator of the distorter is more likely to carry a distorter than an individual with an identical family tree topology, but randomly permuted offspring sexes.

### Clustering Warp results into families

Warp reports the likelihood of carrying a distorter on a per-individual basis. We know, however, that many of the individuals that are likely to carry a distorter are closely related, as clustering of phenotype within families is one of the key ways that Warp classifies individuals as high-likelihood in the first place. To count the unique distorters in the population, we created a method for taking the individual-likelihood results of Warp and grouping high-likelihood individuals together into closely related families. We did this by simple graph traversal. An individual was considered high-likelihood if their likelihood of carrying a distorter was above 40%. From each high-likelihood individual, we walked outward on the graph of relationships, where each individual was a node and each parent-child relationship was an edge. Any high-likelihood individuals within two edges of each other (i.e., a grandparent-child relationship or a sibling relationship) were considered part of the same family. Families could extend further if they consisted of multiple such two-step relationships connected into chains or webs. This algorithm is available in the *relative_clusters* command in github.com/jgbaldwinbrown/tdt.

### Transmission disequilibrium test

In addition to Warp, we also used the transmission disequilibrium test (Spielman et al. 1993) (TDT) to test pedigrees for distortion. We included trios in the TDT in a manner consistent with the expected pattern of inheritance of a *Y*-linked distorter. A valid trio consists of a potential distorter-carrying father, a mother of any genotype, and a child. Because we are searching for *Y*-linked distorters, any male offspring in such a trio carry the same putatively distorting *Y*-chromosome, while female offspring do not. We identified all such trios descended from the (presumed) distorter-carrying progenitor of the pedigree. Then, in keeping with the TDT, we performed a chi-squared test of the expectation of equal numbers of male and female progeny with one degree of freedom. A low p-value here indicates rejection of the null hypothesis and supports the presence of a distorter. Confidence intervals for the TDT were calculated in R using qbinom() with p = 0.995 and p = 0.005, n = family size, and prob = 0.5.

## Key resources table

| REAGENT or RESOURCE | SOURCE | IDENTIFIER |
| --- | --- | --- |
| Deposited data |  |  |
| Software and algorithms |  |  |
| Assorted scripts | This paper | https://github.com/jgbaldwinbrown/jgbutils |
| Transmission disequilibrium test | This paper | https://github.com/jgbaldwinbrown/tdt |

## Supplementary Materials

## Example Bayesian calculations used by WARP

Assume the allele D is dominant, makes all carriers affected (100% penetrant), and has an allele frequency of 0.01. Here, *M* and *F* refer to affected (i.e., male) and unaffected (i.e., female).

Bayes’s theorem applied to probabilities of parental genotypes:

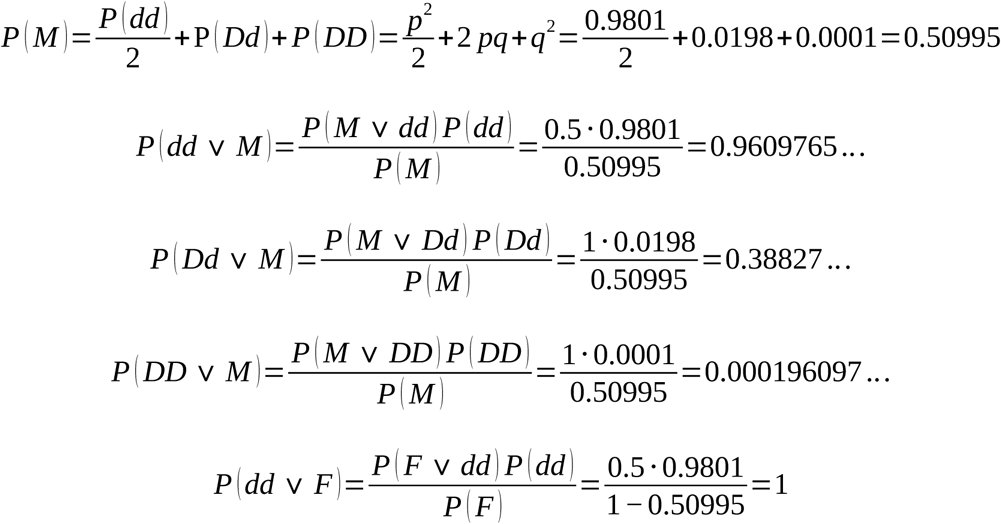

Thus, the probabilities of genotypes of offspring are as below. Bayes’ rule can be further applied to update these probabilities if the sex of the offspring is known. Here, the father’s phenotype will be marked fF or fM, and the mother’s will be marked mF or mM. Additionally, the event of the two parents having opposite affected statuses, equivalent to *fF ∩mM ∪ fM ∩ mF*, will be notated as H.

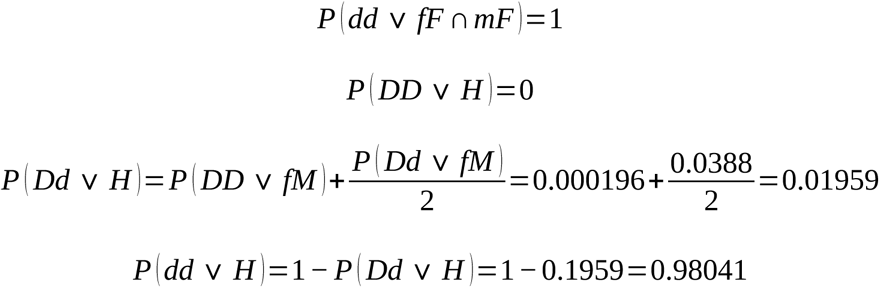

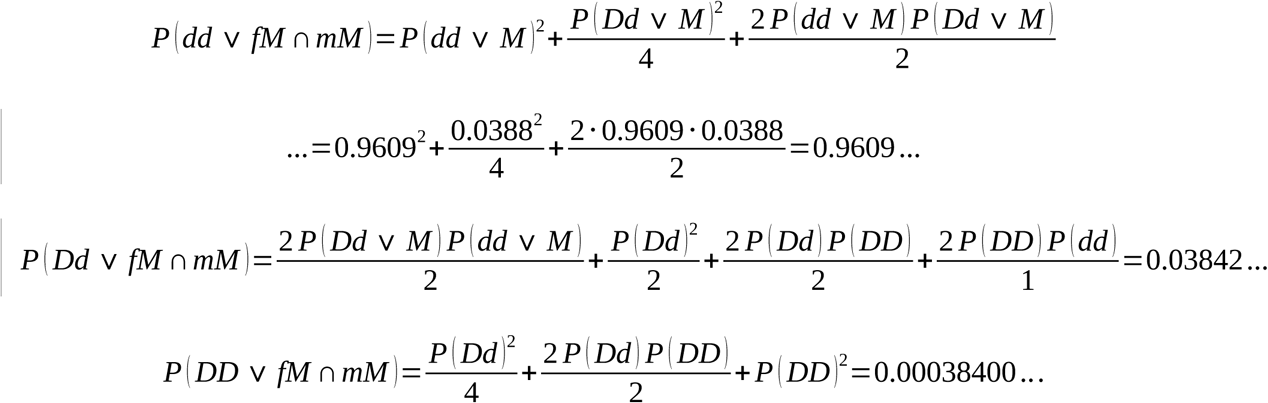

## Supplementary figures

**Supplementary Figure 1.**
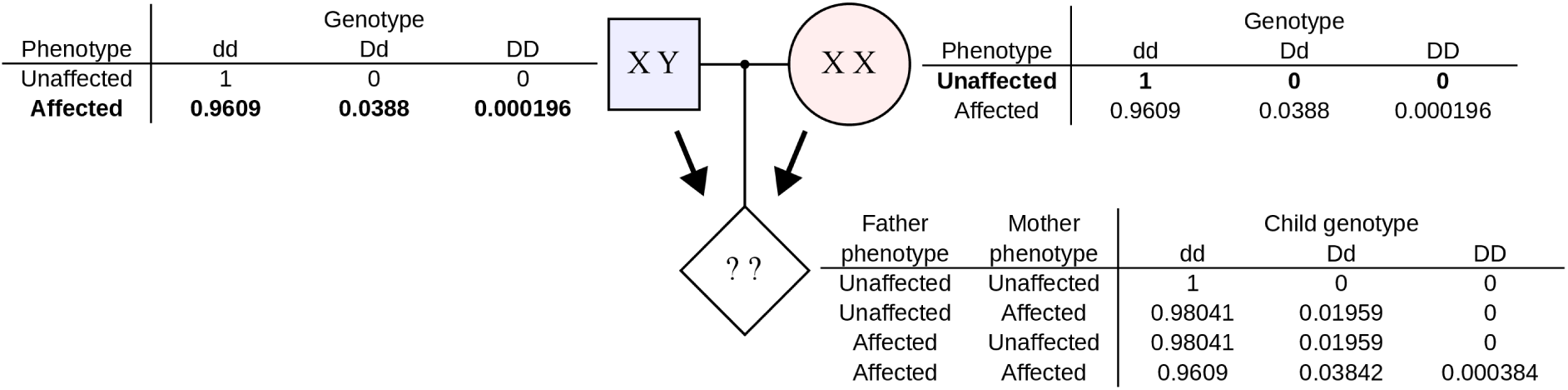
WARP has high power to infer the likelihood of carrying a distorter in pedigrees regardless of penetrance, inheritance pattern, and other challenges. WARP propagates information about the likelihood of carrying a distorter from parent to child, and from child to parent, using Bayesian network propagation (a combination of Bayes’ rule and the chain rule). Here, we depict the probability of two parents transmitting a dominant distorter, D, to a child, assuming that the Dd and DD genotypes cause the affected status 100% of the time, the dd genotype causes the affected status 50% of the time, and the population allele frequency for D is 0.01. The parental genotype likelihoods are calculated using Bayes’ rule, and the child’s expected genotypes are calculated using the chain rule. See Supplementary Materials for detail. In our study, these assumptions would be used when looking for a Y-chromosome distorter with a 100% distortion ratio and an allele frequency of 0.01.

**Supplementary Figure 2.**
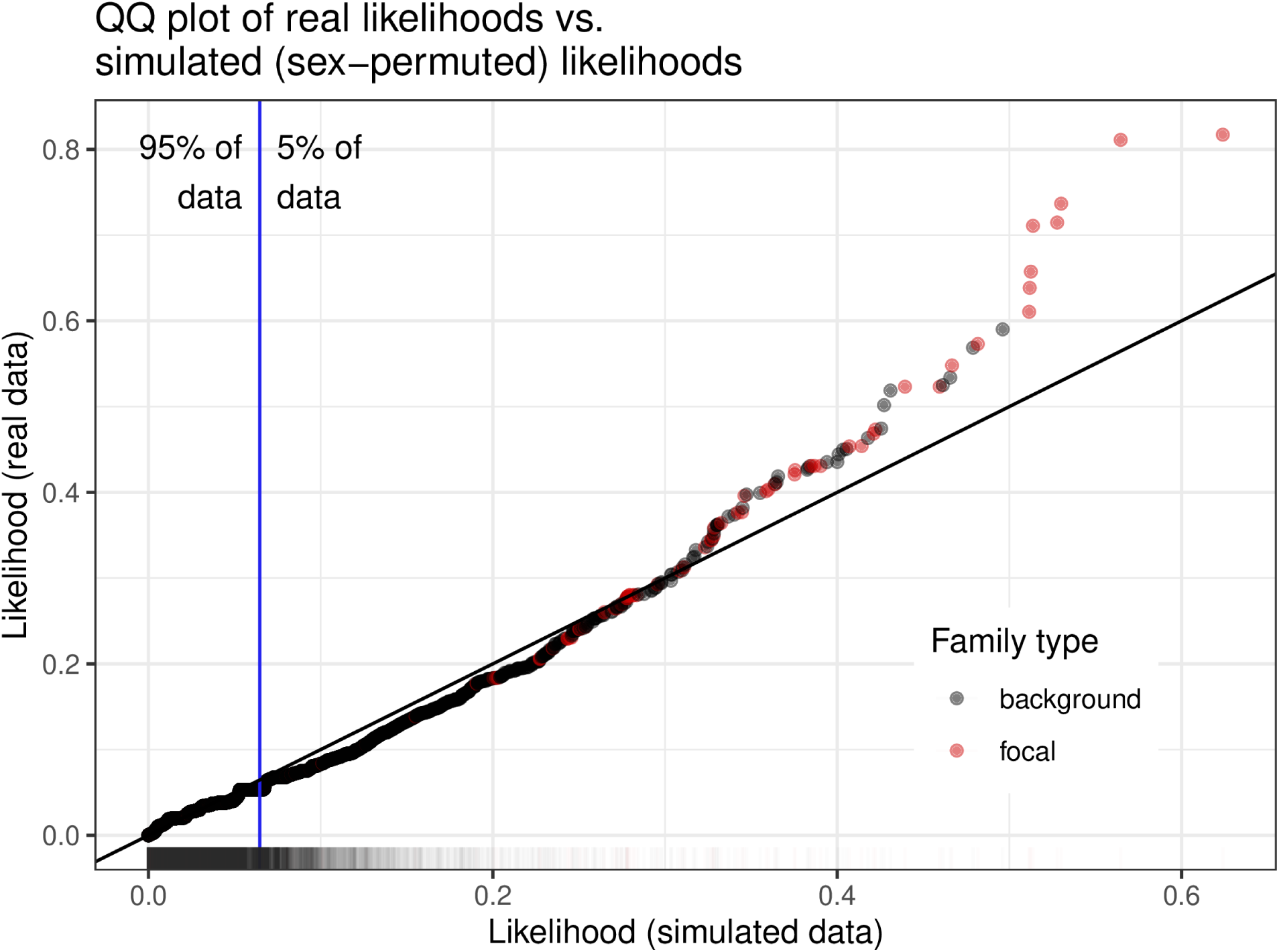
Likelihoods for highly Y-distorted individuals lie outside the distribution of simulated individuals. This Q-Q plot shows the distribution of Bayesian likelihoods of carrying a Y-biased distorter in the true data (Y axis) and in simulated data (X axis, see Methods for details). The black line is the line of 1:1 correspondence, and the blue line represents the 95^th^ percentile (95% of all data is left of this line). There is a notable deviation away from the 1:1 line in a small number of individuals, most of which are in the putative Y-distorting family (family cluster 2, red).

**Supplementary figure 3.**
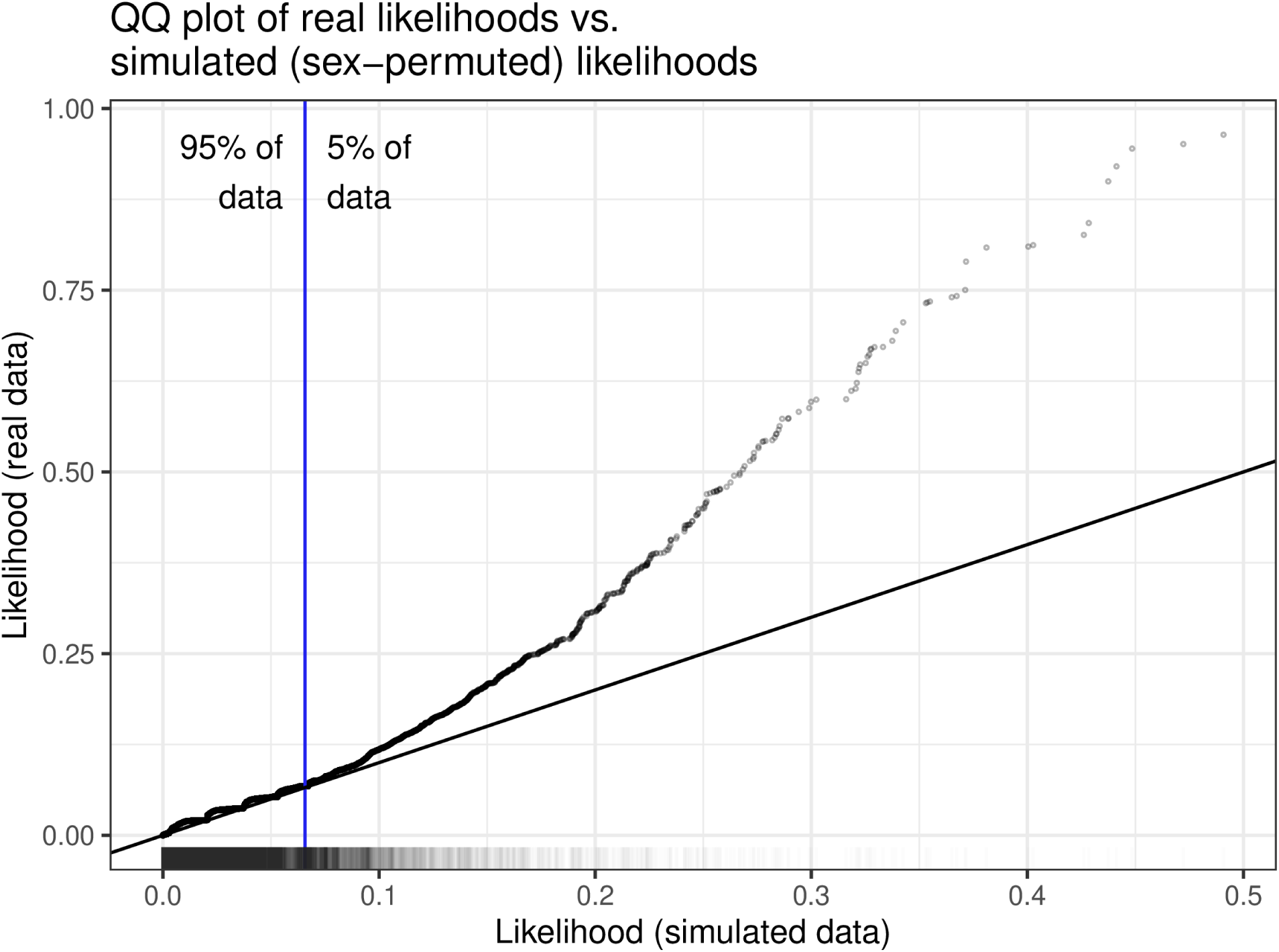
Likelihoods for highly X-distorted individuals lie outside the distribution of simulated individuals. This Q-Q plot shows the distribution of Bayesian likelihoods of carrying an X-biased distorter in the true data (Y axis) and in simulated data (X axis, see Methods for details). The black line is the line of 1:1 correspondence, and the blue line represents the 95^th^ percentile (95% of all data is left of this line). There is a notable deviation away from the 1:1 line in a small number of individuals.

